# Gene expression studies of *WT1* mutant Wilms tumor cell lines in the frame work of published kidney development data reveals their early stem cell origin

**DOI:** 10.1101/2022.06.10.495603

**Authors:** Brigitte Royer-Pokora, Wasco Wruck, Manfred Beier

**Affiliations:** Institute of Human Genetics, Medical Faculty, Heinrich Heine University Düsseldorf, Germany; Institute for Stem Cell Research and Regenerative Medicine, Medical Faculty, Heinrich Heine University Düsseldorf, Germany

## Abstract

In order to get a better insight into the timing of *WT1* mutant Wilms tumor development, we compared the gene expression profiles of nine established *WT1* mutant Wilms tumor cell lines with published data from different kidney cell types during development. Publications describing genes expressed in nephrogenic precursor cells, ureteric bud cells, more mature nephrogenic epithelial cells and interstitial cell types were used. These studies uncovered that the *WT1* mutant Wilms tumor cells lines express genes from the earliest nephrogenic progenitor cells, as well as from more differentiated nephron cells with the highest expression from the stromal/interstitial compartment. The expression of genes from all cell compartments points to an early developmental origin of the tumor in a common stem cell. Although variability of the expression of specific genes was evident between the cell lines the overall expression pattern was very similar. This is likely dependent on their different genetic backgrounds with distinct *WT1* mutations and the absence/presence of mutant *CTNNB1*.

## Introduction

The kidney develops through reciprocal interactions between the stromal/interstitial, mesenchymal and ureteric lineages. The correct signalling between these lineages maintains a balance between self-renewal and induction of differentiation [1]. The ureteric bud (UB) invades the metanephric mesenchyme (MM) and MM cells condense around the UB tip to form the cap mesenchyme (CM). The CM contains the nephrogenic progenitor cells (NPC) that give rise to all epithelial cells of the nephron. The NPC undergo mesenchymal to epithelial transition to form the successive structures: pretubular aggregates (PTA), renal vesicles (RV) and comma- and S-shaped bodies (CSB, SSB). Through the capillary loop stage, the mature and functional glomerular and tubular structures are formed. The UB forms the cells of the collecting duct system (CD). In the mouse, genes expressed in early kidney precursor cells are *Osr1, Eya1, Pax2, Wt1, Six1, Six2, Hox11* paralogs, *Cited1* and *Gdnf*. *Osr1* is expressed in the earliest kidney stem cells that give rise to the majority of all cell types in the kidney [2]. Osr1 function is required for NP cell survival and together with Six2 regulates the balance between cell renewal and nephron differentiation [3]. *Eya1* is expressed in mesenchyme and deletion leads to renal agenesis [4]. Eya1 probably acts at the top of the genetic hierarchy to control kidney organogenesis and acts as critical regulator for *Six1*, *Pax2* and *Gdnf* in a molecular pathway responsible for specification of the metanephric mesenchyme [5]. *Eya1, Six2* and *Pax2* are coexpressed in renal progenitors. Through studies of *Pax2* knock out mice it was found that *Pax2* functions to maintain NP cells, to repress the renal interstitial program and to maintain the lineage boundary between nephron and interstitial cells [6]. *Six2* is autonomously needed for self-renewal and maintaining nephron progenitor cells [7,8].

Uninduced *Six2^+^* CM is characterized by *Cited1* expression and *Cited1* is turned off when the cells are induced [9,10]. In the human kidney *CITED1* is continuously expressed in more differentiated cells of the PTA, RV and SSB [11]. Transient activation of canonical Wnt signalling in *Six2^+^* progenitor cells induce mesenchymal epithelial transition (MET) and tubulogenesis. A regulatory complex of Six2 and Lef/Tcf factors promotes progenitor maintenance. When β-Catenin enters this complex nephrogenesis is initiated [12,13]. Therefore, the balance between Six2-dependent self-renewal and canonical Wnt signalling regulates nephrogenesis. In addition, it was described that the synergistic action of *Osr1* and *Six2* functions to antagonize Wnt directed nephrogenic differentiation and to maintain nephron progenitors [3].

In human kidney development *SIX1* was identified as a *SIX2* target and overlapping expression of both genes was observed in fetal nephron progenitors [14]. ChIP-seq identified a number of potential target genes and the Six2 binding sites mapped to enhancers of genes associated with kidney function [14]. Their study also showed that in human kidney SIX1 and SIX2 bind the same DNA binding motif and among the target genes are *SIX1, SIX2, WT1*, *EYA1* and *OSR1,* but each can form independent regulatory complexes. Both factors, SIX1 and SIX2 bind to their own enhancers to mediate transcriptional activation and they cross-regulate *SIX2* and *SIX1* genes, respectively [14]. One of the most highly regulated targets of SIX2 was *SIX1*. *OSR1* shows comparable expression levels with *SIX1* in human and mouse [14]. In mouse development, *Six1* is expressed in pluripotent renal epithelial stem cells and after the first round of branching it is no longer detected, whereas in human kidney SIX1 activity is found beyond the initial round of branching [14,15].

The earliest expression of *Wt1* in the kidney was identified in the intermediate mesoderm and subsequently in metanephric and CM [16,17]. The important role of *Wt1* for normal kidney development was demonstrated by targeted inactivation in embryonic stem cells. The mutation resulted in embryonic lethality and cells of the intermediate mesoderm go into apoptosis and ureteric bud failed to grow out from the Wolffian duct [18]. In the metanephric mesenchyme *Wt1* is required for the production of UB branching signals and for NP cells response to UB-derived nephrogenic signals, demonstrating that this gene is crucial for the cell interactions during kidney formation [18,19]. In later stages *Wt1* is involved in the control of MET and has an essential function in the development and maintenance of podocytes [20–22]. The *WT1* gene is also involved in the development of a subtype of Wilms tumors, where inactivation/mutation of both alleles was identified [23,24]. These tumors also often harbour additional mutations in *CTNNB1* resulting in activated Wnt signalling [25–28]. This Wilms tumor subtype has a stromal-predominant or stromal-type histology and often aberrant mesenchymal cell types such as fat, bone, cartilage and muscle are present [26,29].

Interstitial progenitor cells (IPC) express *Foxd1,* have self-renewal capacity and all interstitial cell types develop from these [30]. The crucial role for interstitial/stromal cells during normal kidney development was demonstrated by targeted disruption of *BF-2* (*Foxd1*) in mice [31]. *Foxd1*^-/-^ mutant kidneys are small and a decrease in nephron numbers is observed with a delay in nephron differentiation [31]. These studies showed that *Foxd1* expressing cells produce signals/factors needed for normal induction of differentiation of the mesenchyme. In the human kidney *FOXD1* is also expressed in *SIX2*^+^ NPC cells, although at a lower level [32]. It was shown that there is a significant divergence between mouse and human renal embryogenesis [11,32]. Specifically, only 27% of mouse IPC marker genes were enriched in human IPC whereas other mouse interstitial maker genes were found to be expressed in human NPC [32].

We have isolated and described thirteen cell lines from patients with *WT1* mutant Wilms tumors [33,34]. These tumor cell lines have no functional wildtype *WT1* and all except two have additional mutations in *CTNNB1*. The timing of loss of wild type WT1 function likely occurs at different time points in patients with somatic and germ line mutations. It is clear that isolated cell lines cannot reflect the complexity that occurs in kidney development, however they can be instructive to visualize the gene expression pattern at specific stages of human kidney development. The aim of this work was to explore the possible developmental origin of the *WT1* mutant subgroup of Wilms tumors by comparing the transcriptomes of the established Wilms tumor cell lines with those identified in early human kidney development.

## Materials and methods

The cell lines and their genetics have been described in detail elsewhere [34]. In brief all have loss of wildtype *WT1* and all except for Wilms3 and Wilms5 have *CTNNB1* mutations. All cell lines were derived from fresh tumor material at the time of surgery either with or without chemotherapy, and Wilms5 was established from a nephrogenic rest. RNA from these nine *WT1* mutant Wilms tumour cell lines were analyzed with Agilent arrays, containing 60mer long oligonucleotides. First, hybridizations were performed in quadruplicates (Wilms1, 2, 3, 4, 5 and 6) with two biological replicates (RNA isolated from different passages) und two technical replicates (hybridization of two arrays with the same RNA). These gave very reproducible results and in all tested cases the results were confirmed with Q-PCR [33]. Therefore, later Wilms8, Wilms10T and 10M were analyzed in duplicates with two biological replicates. The mean of these replicates was determined and used as basis for these studies.

### Statistical methods

For the analysis of published data, the gene symbols were extracted from Excel files and the expression of these genes was analyzed with the R statistical environment. To study the expression of genes in the Wilms cell lines from the clusters identified by the different authors a stringent expression level cut off >1000 was used. For some of the heat maps a lower cut off >200 was used as indicated in the Figure legends. Hierarchical clustering of the expression data (i.e., the log-intensities) was based on the Ward method, using Euclidean distance as distance measure.

To determine the difference between the Wilms cells with and without *WT1* expression a p-value was computed with a two-sided t-test for each gene in six group comparisons. The group of *WT1* negative cell lines (Wilms1, 10T and 10M) was analyzed separately versus each of the other *WT1* expressing cell lines (Wilms2, 3, 4, 5, 6 and 8), resulting in six p-values. Genes with p<0.05 were selected and analyzed. For Go term analyses ToppGene was used [35] and genes from the different compartments were compared using Venny and exclusively expressed genes were identified [36].

The Wilms tumor cell line transcriptome was also studied using the normal human fetal kidney Atlas (https://home.physics.leidenuniv.nl/~semrau/humanfetalkidneyatlas/). This was done exemplary for one cell line, Wilms8, and after removal of all ribosomal genes, genes with an expression >10 000 were studied for their expression in kidney using the algorithm “sets”. As not all of these genes are expressed in normal fetal kidney, genes with an expression of a Z-score >2 were manually selected, resulting in 550 genes. These were then used to create heat maps and their allocation to the different compartments was analyzed and the number of expressed genes in normal kidney was determined.

### Gene set variation analysis with respect to developmental kidney gene sets

The mean values of the normalized gene expression data were filtered for genes which had expression signals greater than a threshold of t=200 in all experiments. These data were transformed to a logarithmic scale (base 2) and filtered for the upper quartile of the coefficient of variation to find genes with the highest variation for the subsequent Gene set variation analysis (GSVA) [37]. We retrieved the database of developmental kidney gene sets and these were tested against the single cell sequencing clusters detected in two publications [32,38]. The gene expression data filtered as described above were subjected to the gsva function from the R/Bioconductor [39] GSVA package employing these single cell cluster gene sets. The heatmap of the resulting GSVA enrichment scores was drawn with the function heatmap.2 from R gplots [40] package using the “matlab.like” colour palette.

According to the GSVA clustering the samples were assigned to two groups with differential up- and down-regulation of cell cycle. Group1 “CCdown” contained samples Wilms1, Wilms4, Wilms5, Wilms6 and Wilms10M, while group2 “CCup” contained samples Wilms2, Wilms3, Wilms8 and Wilms10T. To determine the statistics of the resulting two groups, genes were filtered with expression >200, limma-p-value <0.05 and a ratio >1.33 between CCup/CCdown and a ratio <0.75 for down-regulation. These were used to make Semrau heatmaps to characterize the CCup stages NPCd/PTA/IPC and CCdown SSBm/d, DTLH, UCBD stages.

## Results and discussion

A comparison between the individual cell lines revealed that they are very similar irrespective of their different *WT1* and *CTNNB1* mutations (Fig 1) To explore whether their gene expression profiles correspond to specific kidney compartments we used published RNA sequencing data sets from human kidney development. In our studies Agilent arrays containing long oligonucleotides from human genes were used. Thus, the data sets using these different technologies cannot simply be compared directly. Therefore, we analyzed the published data by asking which compartment or developmental stage specific genes from the human data sets were also expressed in the Wilms cells and analyzed their expression level.

**Fig 1.**
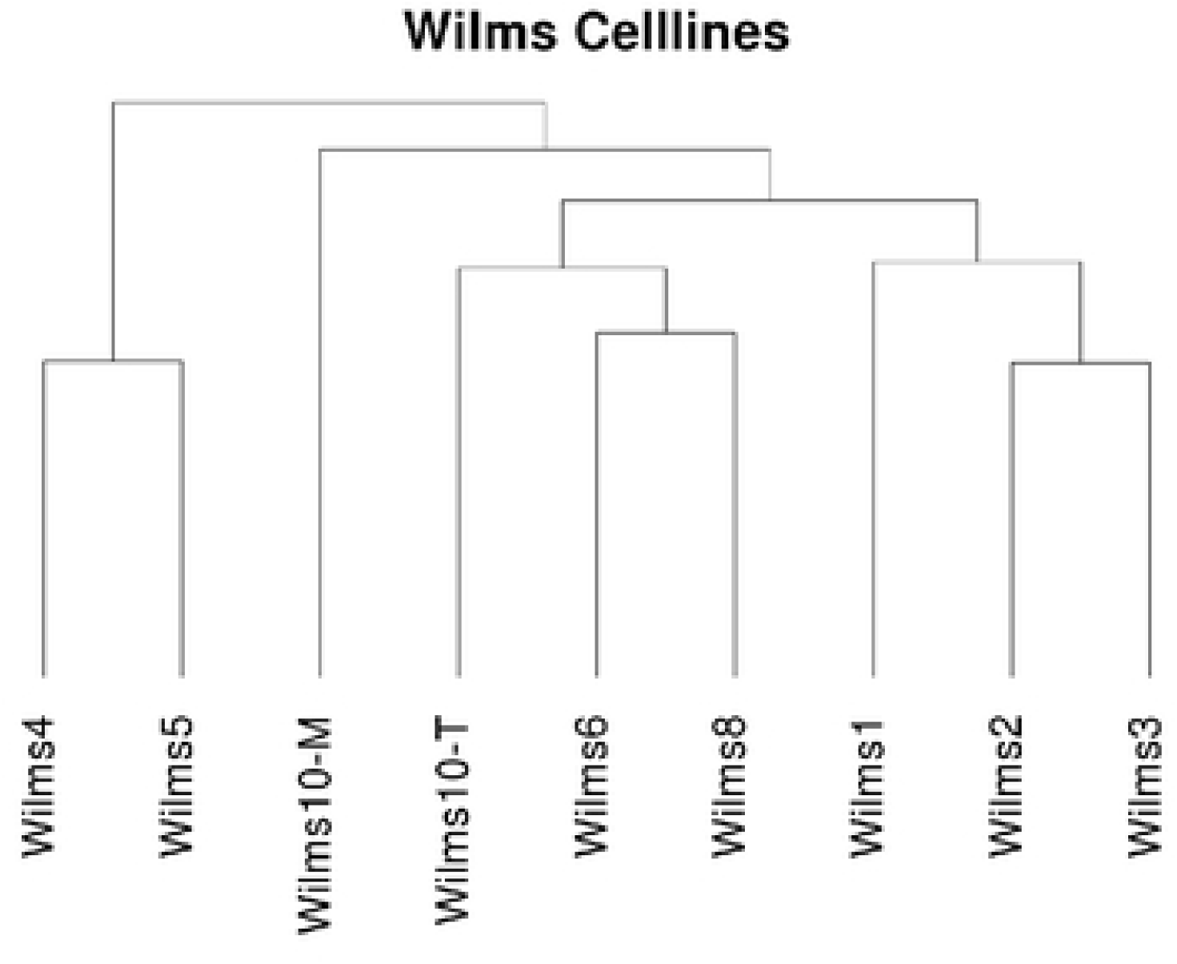
Clustering of all Wilms cell lines demonstrating their close relationship. Cluster method: ward

### Comparison of Wilms tumor cell line transcription profiles with human kidney gene expression

For this purpose, several publications were used that have identified different clusters during human fetal kidney development [15,32,38]. The publications describe genes that are enriched or differentially expressed in the various kidney cell types. These were called clusters by the authors and the studies come up with different gene sets with some overlap between the different authors clusters. Therefore, we analysed the expression of genes from the clusters in the Wilms cell lines from each publication separately.

Lindström et al., showed that human kidney development differs from that in the mouse in various aspects [11,32]. Their work showed that many IPC-associated regulatory factors are also active in NPC and that NPC-associated factors remain active in differentiating nephrons. For example, *FOXD1, MEIS1* and *SIX2* are expressed in IPC and NPC although at a lower level in the latter. These genes are also co-expressed in Wilms cell lines.

These authors used a technique called MARIS (method for analysing RNA following intracellular sorting) to isolate human *SIX2^+^/MEIS1^+^* (NPC) and *SIX2^-^/MEIS1^+^* (IPC) cells for RNA-Seq studies. They showed that with this technique a higher expression of nephron progenitor markers can be observed in the NPC cells as compared to *ITGA8^+^* enriched NPC [14,32] and a lower expression of genes expressed in differentiating nephrons.

We first evaluated the genes that Lindström et al., have identified as differentially expressed between human NPC and IPC cells. Of the 534 genes with enriched expression in NPC, 207 (38.8%) are also expressed in Wilms cells (cut off > 1000) (Fig 2A). These genes were analyzed with ToppGene and the most significant Go term molecular function is “cell adhesion molecule binding” with 18 genes (*P*=3.46E-6) and genes with a high expression of >1000 are listed Fig 2B. The second Go term is “signalling receptor binding” with 37 genes (*P*=8.02E-5) and the highest expressed genes are listed. Another significant term was “actin binding” with 14 genes (*P*=3.30E-4) (Fig 2B). When the same analysis was performed with all 534 normal human NPC enriched genes, “actin binding” is the most significant Go term molecular function with 31 genes (*P*=7.05E-7). The term identified in Wilms cell lines as most significant “cell adhesion molecule binding” is second (*P*= 1.11E-5) (Fig 2C) and the highly expressed genes (transcript per million, TPM>10) are listed. The third term is “signalling receptor binding” with 75 genes (*P*=2.54E-5) and here also the highest expressed genes in normal NPC cells are listed. As most of the highly expressed genes in normal NPC and Wilms cell lines are identical this indicates that the Wilms cell lines have very similar but slightly distinct characteristics as normal human NPC cells. Some important NPC enriched genes that are not expressed in the Wilms tumor cell lines are *CITED1, COL9A2, EYA1, DACH1, ECEL1, HES6, HEY1, LYPD1, NNAT, PAX2, PCDH15, RSPO1, SLIT1* and wild type *WT1*.

**Fig 2.**
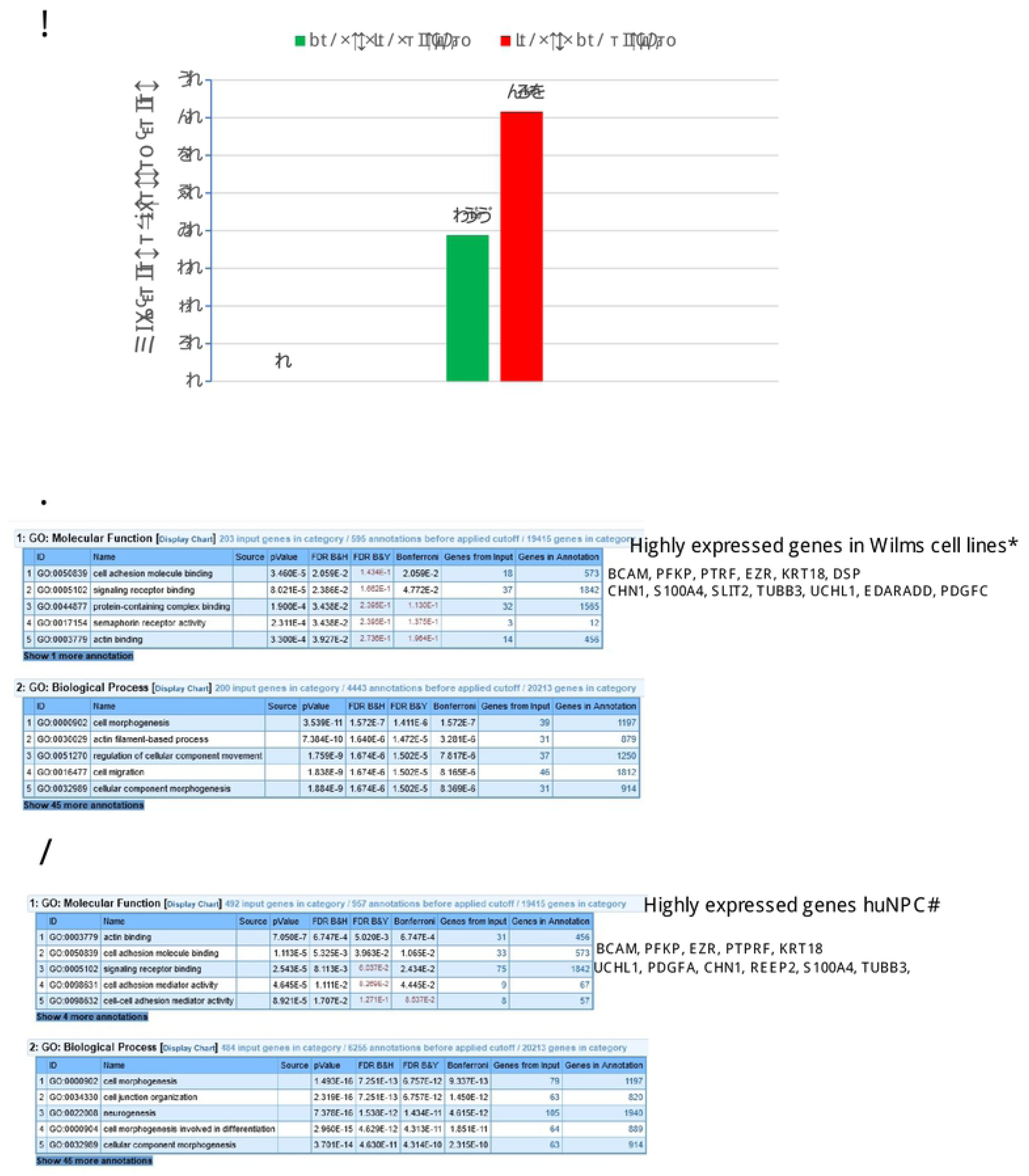
Analysis of genes from the enriched NPC compartment expressed in Wilms cells [32]. **A)** Percent of genes from the Lindström NPC enriched and IPC enriched compartments expressed in Wilms cell lines (>1000). **B)** ToppGene analyses of the 207 NPC enriched genes expressed in common in Wilms cells. *: genes, with an expression level between 3000 and 260 000 **C)** ToppGene analysis of all 534 normal huNPC enriched genes [32]. #: genes with a mean TPM (transcript per million), between 10 and 172 [32].

The analysis of IPC enriched genes uncovered that 71.6% of the genes are also expressed in Wilms cells >1000 (Fig 2A). The most significant Go term molecular function is “extracellular matrix structural constituent” with 34 genes (*P*=1.18E-24), and genes are listed according to their expression level in Wilms cells (Fig 3A). The same term is the top term for normal IPC genes identified by Lindström et al., and genes are listed according to their expression level. All significant terms are identical in normal IPC and Wilms cells but with a different order (Fig 3B). These results support the notion that the *WT1* mutant Wilms tumor cells are more related to interstitial cells although they are not identical and have adopted an aberrant path as they also robustly express genes from NPC population as well.

**Fig 3.**
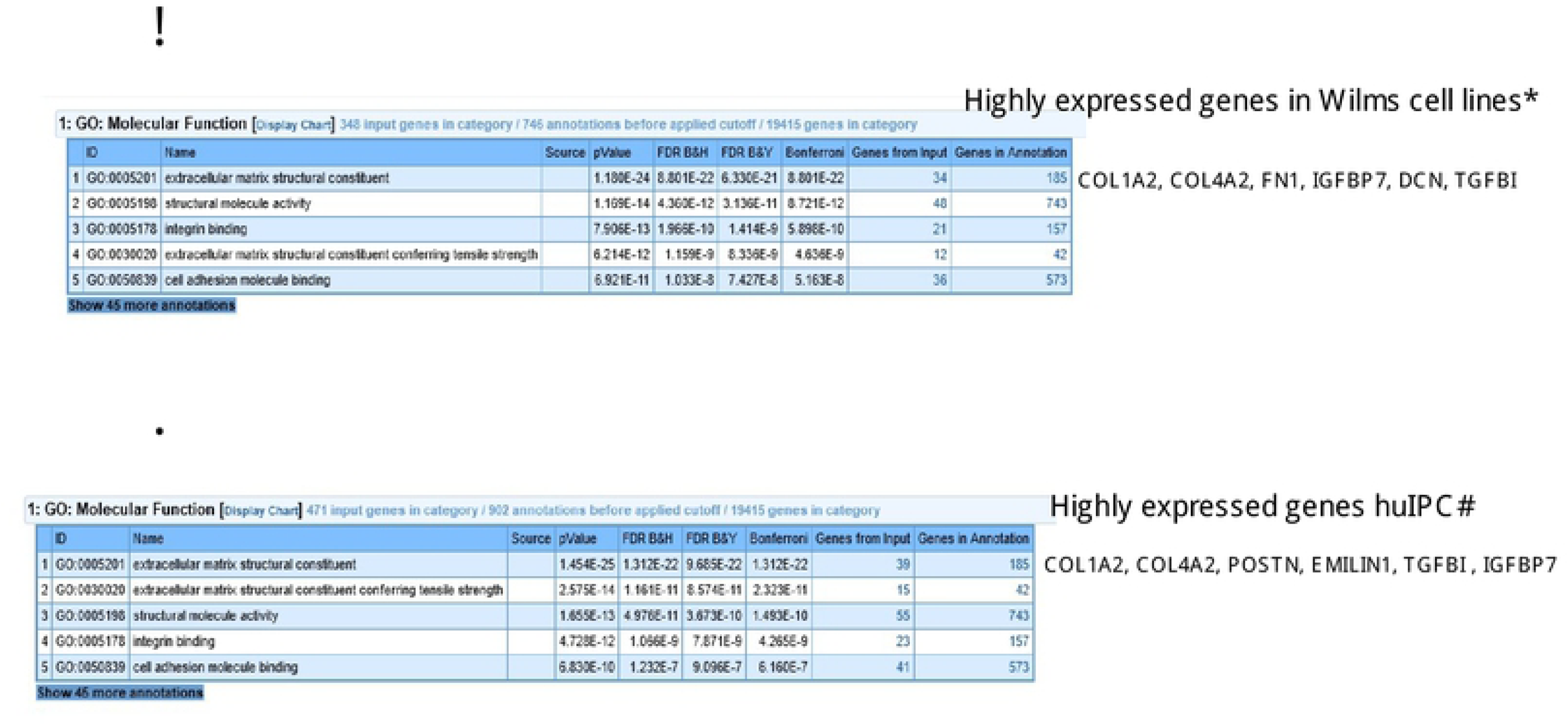
Functional analysis of genes of the IPC enriched genes [32]. **A)** ToppGene analyses of the 360 IPC enriched genes also expressed in Wilms cells >1000. *: genes with an expressed level between 20 000-270 000 in Wilms cell lines. **B)** ToppGene analysis of all 503 normal huIPC enriched genes [32]. #: genes with a mean TPM between 70 and 281 [32].

To study the cellular diversity of human nephrogenic cells in more detail the authors performed single cell RNA-Seq on 2750 predominantly mesenchymal cell types of the cortical nephrogenic niche from 16-week human fetal kidney. Unsupervised clustering uncovered 12 cell populations initially identified by known marker genes and with different numbers of genes. Five of these were from interstitial lineages (cluster 2, 9, 10, 11 and 12), and three from the nephrogenic lineage (cluster 4, 5 and 6). In addition, one cell population corresponds to vascular endothelial cells (cluster 3), two to proliferating cells (cluster 7 and 8) and one to immune cells (cluster1). We analyzed how many genes from each cluster are expressed in the Wilms cell lines (>1000) and the highest number of genes is found for the developing vascular cell cluster 3 (678/933, 88,3%) and from proliferating clusters 7 (721/890, 80.9%) and 8 (707/860, 82.2%) (Fig 4A). The high number of genes expressed from the proliferating clusters is not surprising as tumor cells have a high proliferative potential. The lowest number of genes expressed in Wilms cells is from cluster 6, corresponding to differentiating nephron cells (551/960, 57%). This demonstrates that the cell lines do not differentiate in culture. S1 Table shows the key genes for each cluster, the number of genes in each cluster, as well as the highest expressed genes in the Wilms cell lines.

**Fig 4.**
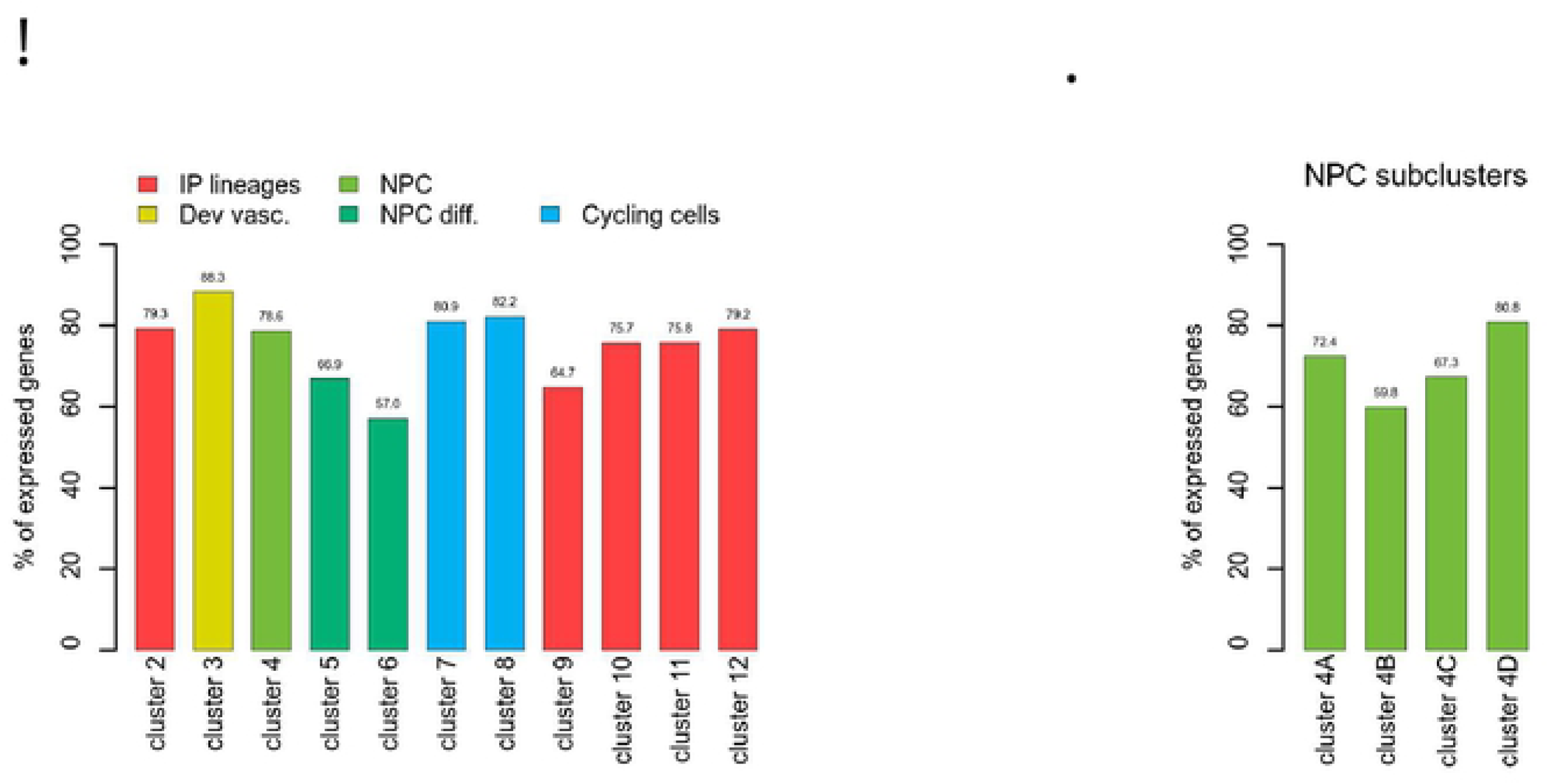
Analysis of expressed genes from the Lindström clusters 2-12 in Wilms cells. **A**) The percentage of genes expressed in each cluster are shown. Cluster 1, immune cells, was not analyzed. The color coding shows which clusters are part of the NP (green) or IP lineages (red). The differentiating NP cells, cluster 5 and 6 are darker green. Two clusters of cycling cells are turquois and developing vasculature cluster is yellow. **B)** Exclusively expressed genes from the 4 NPC subclusters were analyzed and the percent of expressed genes in Wilms cell lines are shown, cut off >1000.

The authors further studied the heterogeneity of NPC cells and identified 4 subclusters, called A for self-renewing cells, B or primed NPC, C for differentiating cells and D for proliferating cells. We analyzed which genes are exclusive for each cluster (S1 Fig) and how many of these from each subcluster are expressed in the Wilms cell lines (>1000) (Fig 4B). Here the highest percentage of genes are expressed from the proliferating cluster 4D (80.8%), as would be expected for fast growing tumor cells. These analyses showed, that a large number of NPC specific genes are robustly expressed in Wilms cells.

We also studied genes from clusters 9, 10, 11 and 12 that are exclusively expressed in the Wilms cell lines. A summary of all analyses is shown in Fig 5 and the eight highest expressed genes in the Wilms cell lines in each cluster are listed. Genes that are also found in other compartments are shown in black and are underlined. For example, *ANXA2* previously identified as an interstitial marker gene in the mouse is found in NPC4C and developing vasculature and it is highly expressed in the Wilms cell lines. It has also been assigned to podocytes [38]. Another example is *UCHL1* which is found in cluster 4 and differentiating nephrons (cluster 5) that develop from cluster 4 during differentiation. A full list of genes expressed in the Wilms cell lines from each cluster is found in S2 Table.

**Fig 5.**
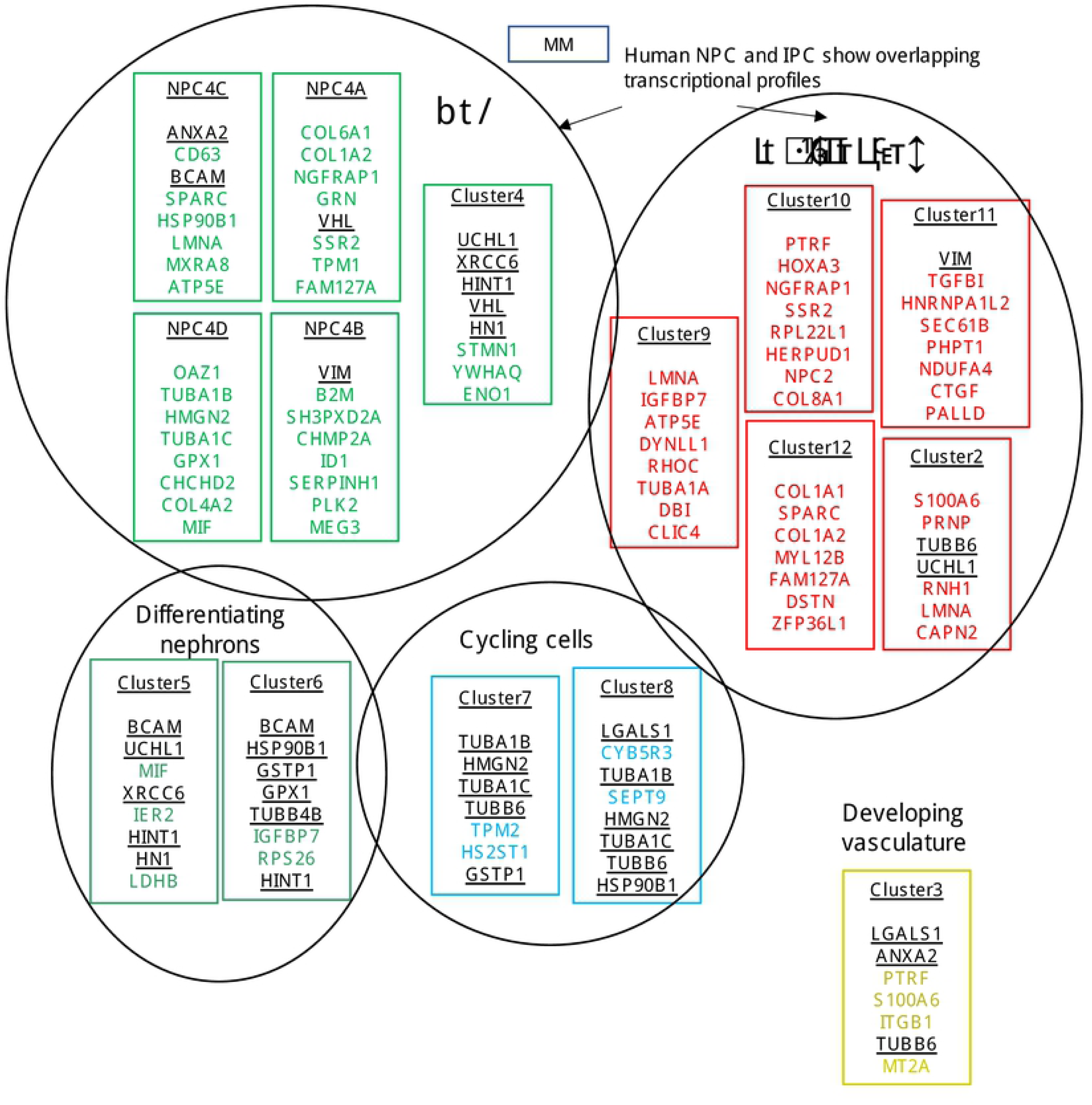
Lindström clusters and expression of genes in Wilms cell lines. First genes that are specific, i.e., only expressed in one cluster were extracted. This was done separately for clusters 4A, 4B, 4C, and 4D from the NPC compartment. Cluster 4 is the parental cluster and is shown in the same circle. The top expressed unique genes in Wilms cell lines are listed and labeled green; black and underlined genes are also listed in other compartments. Clusters from the IPC compartment were also analyzed for genes exclusively expressed in the subclusters, shown in the red circle. The top expressed unique gene in Wilms cell lines are listed and labeled red; black and underlined genes are also listed in other compartments. The top genes for differentiating nephrons are shown in green and for cycling cells in turquois. Cluster1 (immune cells) is not shown.

Hochane et al., identified 22 clusters by single cell transcriptome analysis of 16-week human fetal kidney cells [38]. The clusters were assigned by using the expression of marker genes from the literature of mouse kidney development. The percentage of marker set candidate genes from each cluster is shown in Fig 6A and a list of all genes expressed in Wilms cell lines (<1000) from these clusters are found in S3 Table. The nephrogenic precursor cells (NPC) were represented by four clusters and they express the known markers *SIX2, CITED1, MEOX1* and *EYA1*, with the highest expression in cluster NPCa, the self-renewing NPCs. Of these only *SIX2* is expressed at highly variable levels between the Wilms tumor cells lines. These genes were expressed at a lower level in NPCb, but a higher expression of *GDNF* and *HES1* was observed [38]. In the Wilms cell lines both of these genes are expressed at a medium level. NPCc is characterized by a high expression of *CRABP2*, which is also expressed in the Wilms cells at a medium level. NPCd had a higher expression of *LEF1* and a lower expression of *OSR1, CITED1* and *MEOX1*, whereas the Wilms cell lines express only low levels of *OSR1* and a highly variable level of *LEF1*. The authors noticed that a larger fraction of NPCd cells were in the G2/M phase of the cell cycle and had a high expression of proliferation markers as is observed in the Wilms cell lines. These correspond to the Lindström NPC 4D subcluster.

**Fig 6.**
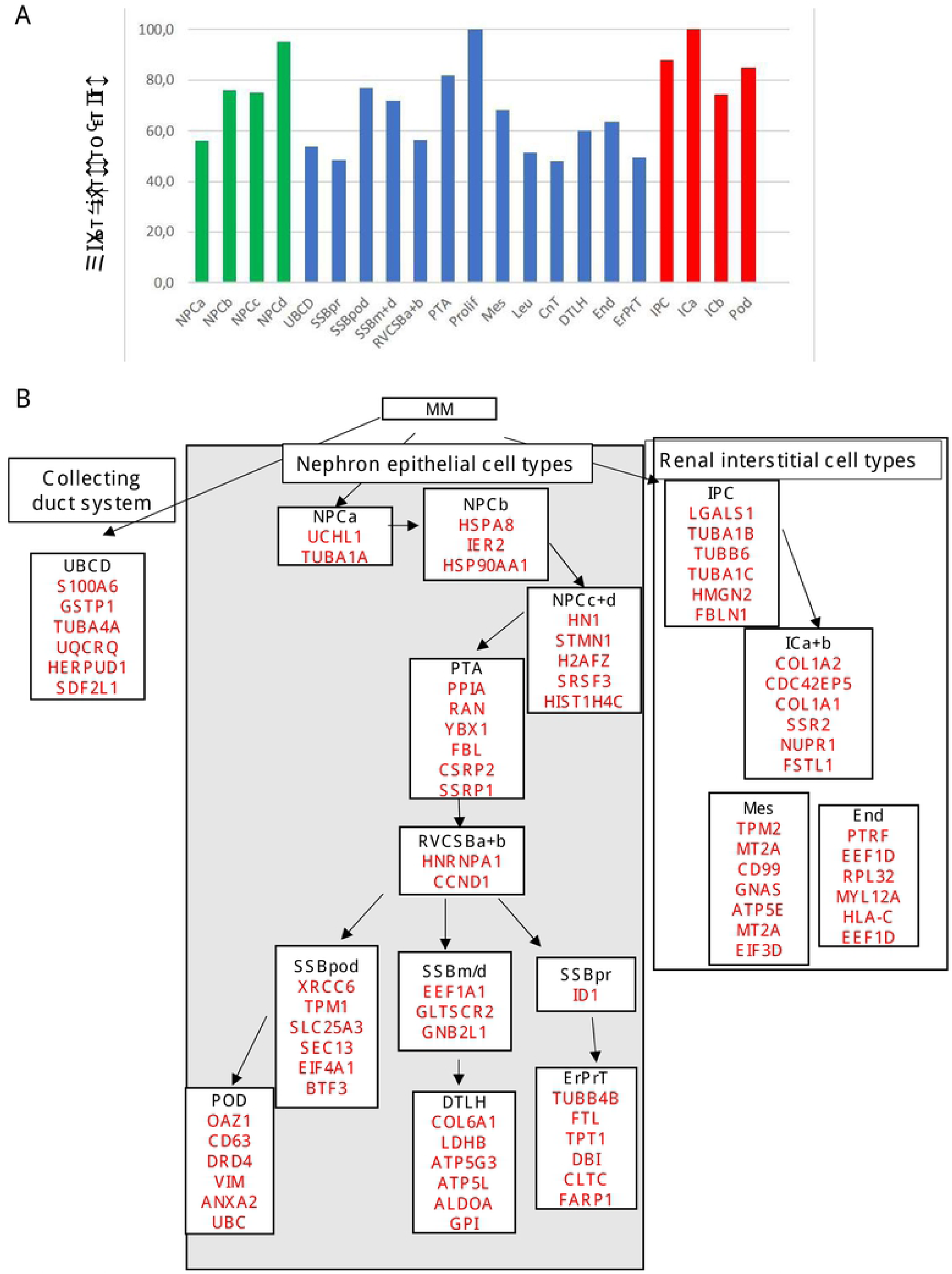
Analysis of expressed genes from the Hochane marker set candidates in Wilms cells. **A**) The percentage of genes expressed in each cluster is shown. **B)** Genes in each cluster from the three linages that are expressed >50 000 are shown. Ribosomal genes were omitted. No genes in the CnT cluster were expressed above 50 000.

When nephrogenesis continues pretubular aggregates (PTA) appear, the precursors of RV and CSB. PTA express a high level of *LHX1*, which is only expressed at a very low level in the Wilms cell lines. A higher expression of the other PTA markers, *JAG1* and *CCND1* is observed in the Wilms cell lines but *WNT4* is not expressed. Several markers distinguish distal and proximal RV, e.g., *SFRP2, DLL1, LHX1* and *CDH6, FOXC2, CLDN1* and *WT1,* respectively. Most of these vary between cells and have a low level, except for a robust expression of the podocyte marker *FOXC2,* and *DLL1* is not expressed. Genes that are expressed in the Wilms cell lines from the next developmental stages are, SSBpr: *CAV2, CDH6* and *AMN,* SSBm/d: *IRX2, POU3F3* and *SIM2*, and SSBpod: *XRCC6, FOXC2* and *MAFB*. This is followed by more differentiated cell types, and the genes that are expressed in the early proximal tubule (ErPrT) are *AMN, APOM* and *ANPEP* and from distal loop of Henle (DTHL) are *LDHB* and *HOXD8.* Characteristic genes for CnT that are expressed in the Wilms cells are *SLIT2, BCAT1* and *MEST* and from podocytes many genes are expressed at a very high level, among these are *VIM, ANXA2, ACTG1, PODXL2* and *MAFB.* A high expression of several markers that are characteristic for UBCD is also seen in the Wilms cell lines (Fig 6B). The interstitium is divided into three clusters, IPC, the stem cells and the more differentiated cells types ICa and ICb. Mesangium derived from IPC and endothelial cells correspond to clusters Mes and End. The expression of genes in the Wilms cell lines >50 000 from each cluster is shown in Fig 6B and the expressed genes are seen in S4 Table. Taken together, the Wilms tumor cell lines express markers from the earliest NP and IP stem cells, as well as markers from ureteric bud and collecting duct and more differentiated cell types.

To identify novel markers the authors used two approaches. First, they analyzed for each gene the area under the receiver operating characteristics (AUROC) and combined this with expression filtering. This resulted in 88 marker genes and 11 of these overlapped with 89 literature genes. The expression of these genes in Wilms cell lines is shown in the heat map in Fig 7. All top 4 genes from NPC and IPC domains, as well as from the CnT, UBCD and proliferation clusters are expressed in the Wilms cell lines (>200), but only single genes within the clusters showed a high expression. A lower expression level can be observed for almost all the genes that are characteristic for more differentiated cell types such as RVCSB, SSB, CnT, DTLH, ErPrT and UBCD (Fig 7 and S4 Table).

**Fig 7.**
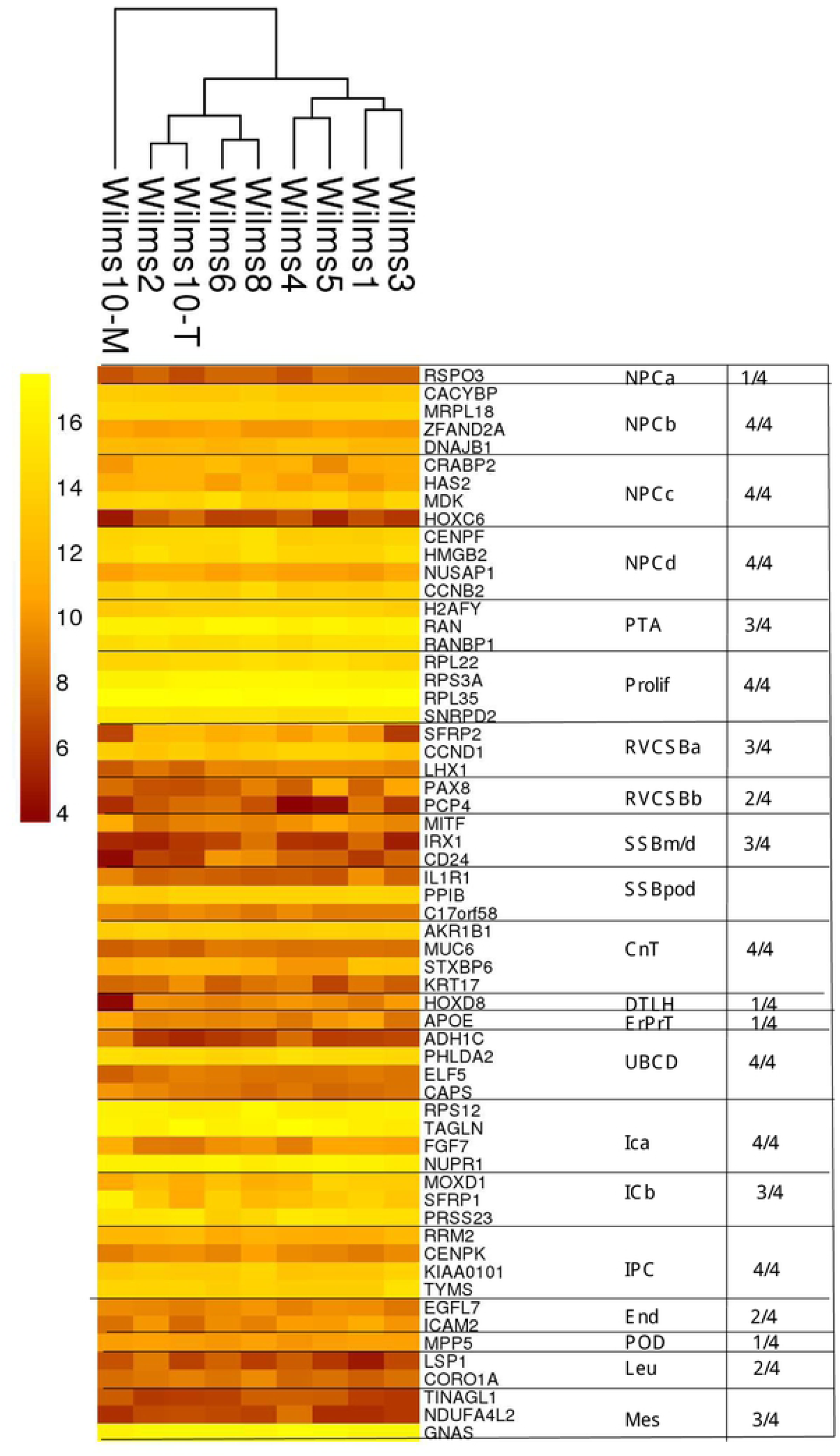
Expression of the TOP 4 genes from the marker set candidates in the Wilms cell lines. In the heat map the expressed genes from the top 4 genes are shown (>200).

As a second method they used the *KeyGenes* algorithm to identify classifier genes among the 500 most highly variable genes. Of the 95 classifier genes 24 were the same as in the marker set and 14 were common in the literature set. The heat maps in Fig 8 and S2 Fig show the expression of *KeyGenes* in the Wilms cell lines. Each compartment has different numbers of *KeyGenes* and all from the NPCd, ICa and IPC compartments are highly expressed, whereas lower numbers and levels of genes from the other compartments are expressed in the Wilms cell lines (S2 Fig). Also, in this analysis it is obvious that genes from SSBm/d, RVCSBa+b are expressed at a much lower level than those from the NPC and IPC compartments (Fig 8 and S2 Fig). Lastly, we analyzed the expression of their literature gene set and here the highest expression level was observed for genes from IPC, ICa+b and mesangium cells, whereas most other genes have a lower expression (S3 Fig). In conclusion, the robust expression of several genes from more mature nephrogenic compartments shows that the pathway for epithelial differentiation is not completely blocked in these cells. It has been observed previously, that individual mouse MM cells at E11.5 express markers of more differentiated nephron cell types [10]. Expression of genes from different lineages is also observed in the Wilms cells, e.g., podocyte markers, *MAFB, PDPN, SYNPO, RHPN1* at highly variable levels between the different Wilms cell lines or a high expression of genes from DTLH, UBCD and ErPrT (Fig 6B). This supports the notion that genes associated with possible future developmental directions can occur in the precursor cells such as Wilms tumor cells.

**Fig 8.**
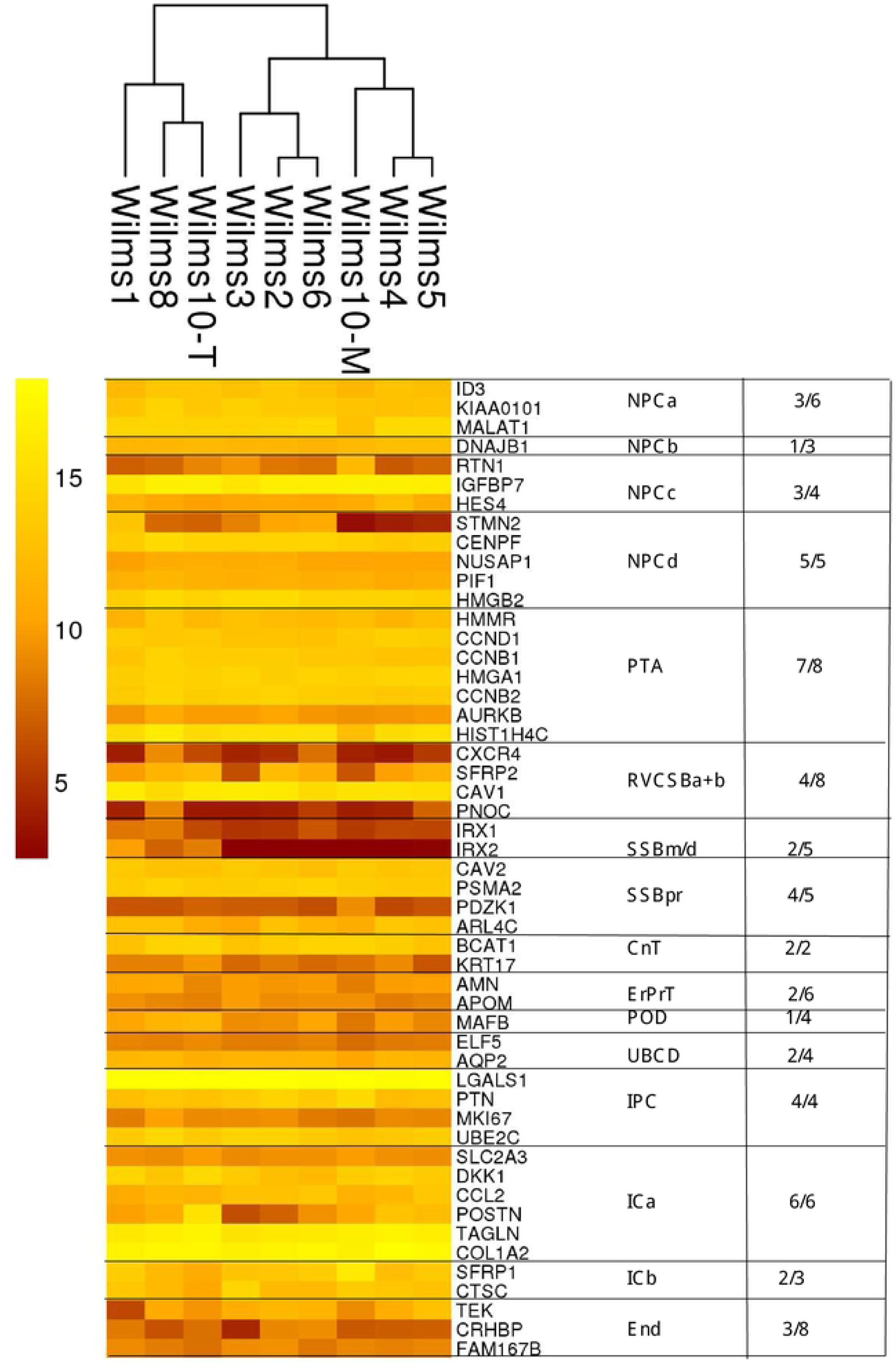
Expression of *KeyGenes* in Wilms cell lines. Each compartment has a different number of *KeyGenes*. The expressed genes from each set are shown (>200) and numbers are listed on the right.

Another way to analyze the Wilms tumor cell line transcriptomes is to study the highest expressed genes for their expression in normal human fetal kidney. Of the 1300 highest expressed genes in Wilms cell lines, 550 genes with an expression Z-score of >2 in normal human kidney were identified. This analysis demonstrated that most of highly expressed genes in Wilms cell lines map to the DTLH compartment, followed by PTA, UBCD, IPC, NPCd and End (S4 Fig). Among the genes with highest Z score, were *MYL7* (NPCb), *PALM2-AKAP2* and *ARHGAP11* (NPCd), *LBX1* (PTA), *GRIND2* (RVCSBb), *CRLF1* (CnT), *NDOR1, ATP1A1* and *COL18A1* (DTLH), *COX6A2* (ErPrT), *FAM65A* (Pod), *CNN1* and *CDKN2C* (IPC), *ACTA2* (Mes) and *TM4SF1* (End). It is interesting that the highly expressed genes in Wilms cells that map to the NPCd and PTA clusters are related to the cell cycle as was observed by Hochane et [38]. As many of the highly expressed gene are allocated to the PTA, DTLH and UBCD compartments, this suggests that a differentiation into these different kidney cell types has been simultaneously initiated in the same cells.

To combine the results from these studies we have established a heatmap of GSVA enrichment scores using the Hochane and Lindström 2018b gene sets with the Wilms tumor samples (Fig 9). The top part of the heat map, shows that Hochane NPCa and NPCd are close to Lindström NPC clusters 4, 4d, 4b and the cell cycles clusters 7 and 8. Hochane IPC maps within these clusters. Here a high score is found in Wilms3, 10T, 8 and 2, whereas Wilms1, 6, 5, 10M and 4 show a lower score. It is interesting that Lindström NPC4A and NPC4C are in the middle of more differentiated cell clusters and among the Lindström interstitial clusters 3, 9, 10, 11, 12 where the Hochane ICb cluster is also found. Hochane IPC, the interstitial stem cells are close to Hochane NPCd. This analysis uncovered that the Wilms cells are separated into two main clusters, where Wilms1, 4, 6, 5 and 10M (group 1, “CCdown”) have a higher score for more differentiated cell type gene sets and a lower score for NPC, IPC and cell cycle gene sets, whereas Wilms3, 10T, 8 and 2 (group2, “CCup”) cell lines have a higher score for NPC and proliferating clusters. The differences between the two groups are most pronounced in the Hochane gene set associated with the NPCd cell type. Hochane et al. described that NPCd is likely more proliferative than the other NPCs and in an intermediate state between the other NPCs and the pretubular aggregate.

**Fig 9.**
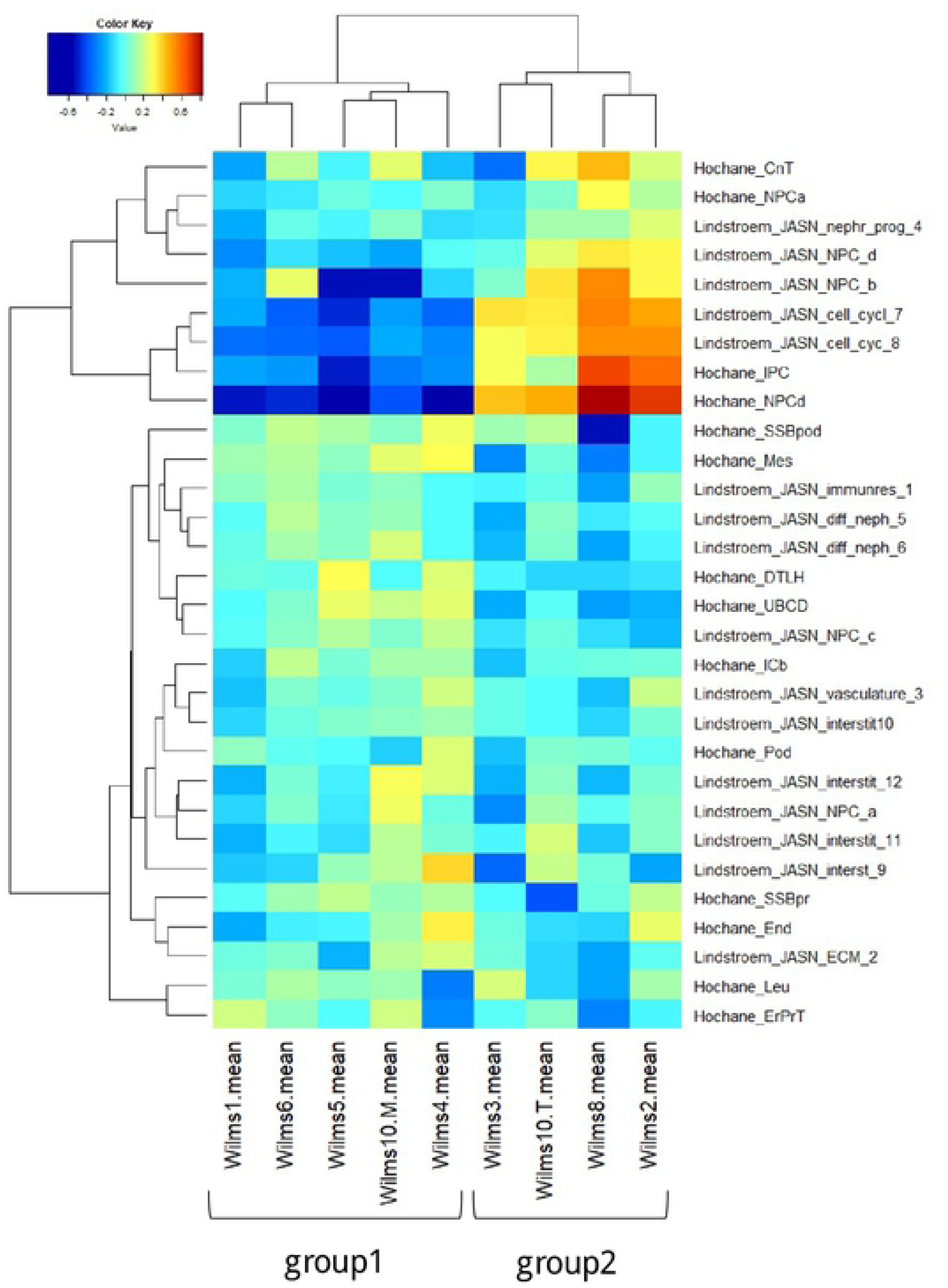
GSVA enrichment scores for the Hochane and Lindström clusters in Wilms cells. Expression values of Wilms tumor samples were subjected to GSVA analysis using gene sets associated with developmental kidney cell types in the single cell sequencing studies [32,38]. In the hierarchical cluster analysis of the GSVA enrichment scores two main clusters of samples emerged: group1 (Wilms Tumor samples 1, 4, 5, 6 and 10M) had higher enrichment in differentiated and lower enrichment in nephron progenitor cell (NPC)-like cells; group2 (Wilms Tumor samples 2, 3, 8 and 10T) had lower enrichment in differentiated and higher enrichment in NPC-like cells.

To explore this result further, we determined the differences between these two groups of Wilms tumor cells lines with a Limma-p-value <0.05, expression <200 and a ratio >1.33 between up/down and a ratio <0.75 for down-regulation. These were used to analyze genes that are expressed in normal human kidney [38] and confirmed the high expression of NPCd, PTA and IPC compartment genes, many of these are related to the cell cycle (S5 Fig). For the group of down regulated genes, expression of genes from more differentiated cell types e.g., UBCD, DTLH and ICb was observed as seen in the GSVA map (S6 Fig).

This was somewhat unexpected as all cell lines proliferate well and show similar features in culture, but the differentiation state of the cells is unknown. We explored which factor might be the reason for this separation and analyzed the preoperative chemotherapy versus primary surgery, the presence or absence of *CTNNB1* mutations, the complete lack of expression of the mutant *WT1* and the time of preoperative chemotherapy in both groups. However, none of these could explain the difference as cells from patients with these differences could be found in both groups. Therefore, the reason for this separation currently cannot be explained and remains to be elucidated in the future.

Another paper describes the study of gene expression in human kidney [15]. We analyzed the expression of genes in Wilms cell lines that correspond to the clusters defined by these authors and the full list of genes from each cluster expressed is found in S5 Table. In S7A Fig the percentage of genes from each cluster expressed in Wilms cells with a cut off >1000 is shown and the number of genes in each compartment is listed in S7B Fig. Here a high percentage of genes from cap mesenchyme (CM), extraglomerular mesangium (EM), loop of Henle (LOH) mesangium (MG) podocytes (PD) and renal interstitium (RI) is seen. S6 Table shows a list of marker genes from these 13 clusters and the eight highest expressed genes in the Wilms cell lines. When the same genes are found in different clusters they are coded with the same color, demonstrating that there is a large overlap of genes between these clusters and therefore these gene sets were not studied further.

Previous studies have suggested that the *WT1* mutant Wilms tumors are closely related to interstitial/stromal cells but here we show that they robustly express genes from the NPC, UB and IP compartments as well as some genes from more differentiated cell types. This further supports the notion that the origin of *WT1* mutant tumors resides in an early kidney stem cell where the underlining mutations result in a deviation from the normal differentiation pathway.

### Individual differences between *WT1* mutant cell lines

Another line of studies was to explore the difference between the cell lines with and without *WT1* expression. Although all cell lines have only mutant *WT1* DNA alleles the RNA expression differs between the cell lines [41]. Wilms1 has an almost absent expression of *WT1* RNA and a homozygous *WT1* deletion is present in Wilms10T and 10M. These three cell lines were analyzed as a group and compared against the other cell lines. A p-value of <0.05 in all six t-tests was found for 114 genes, with a higher expression of 49 and a lower expression of 65 genes in cells with lack of *WT1* expression (Fig 10A and B, respectively). Two genes, *SIX1* and *SIX2* that are normally expressed in nephron progenitor cells show a lower expression in cells without *WT1* (Fig 10B). This correlated with a higher expression of *OSR1* and *OSR2*, but these did not reach a p*-*value of <0.05.

**Fig 10.**
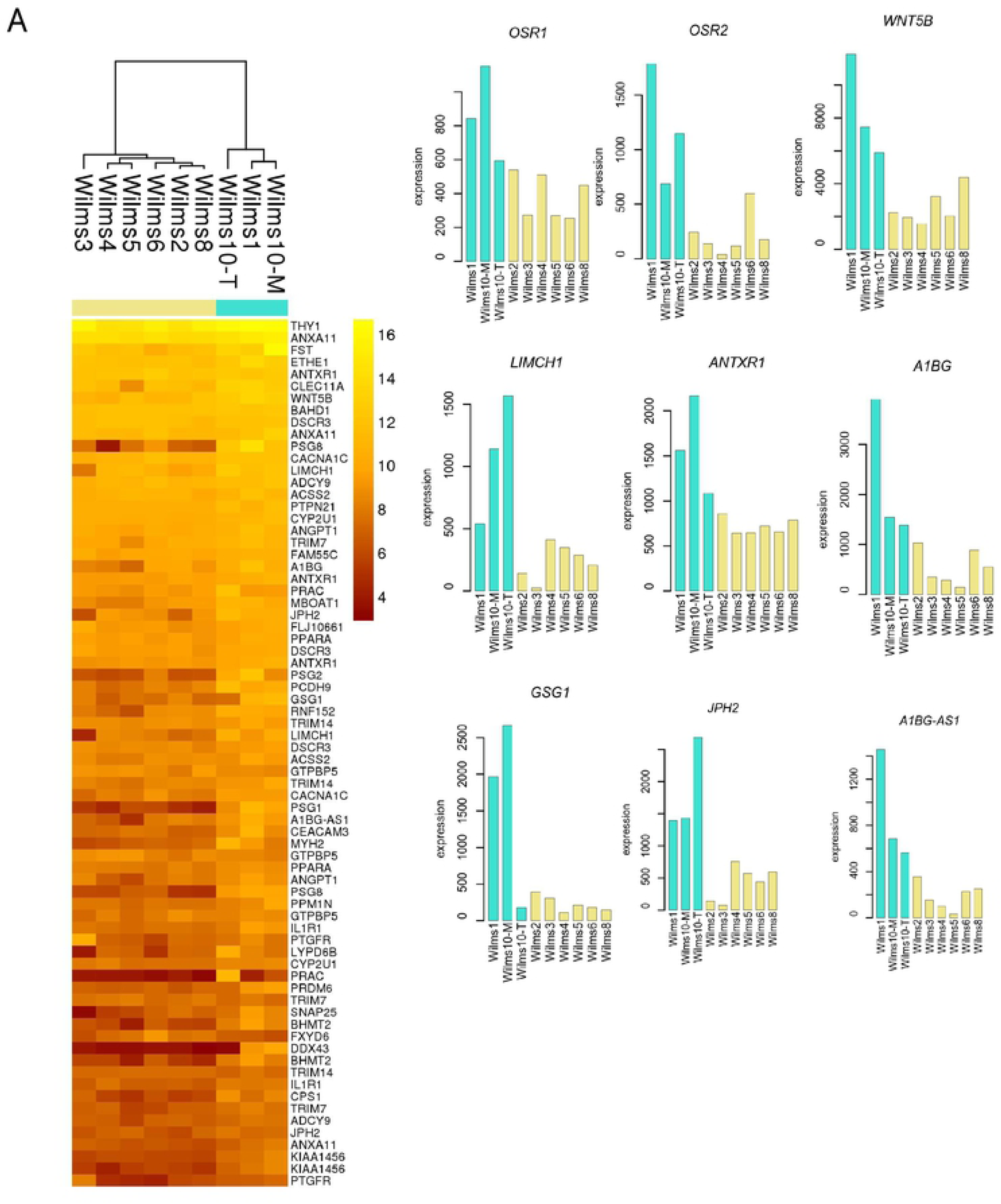
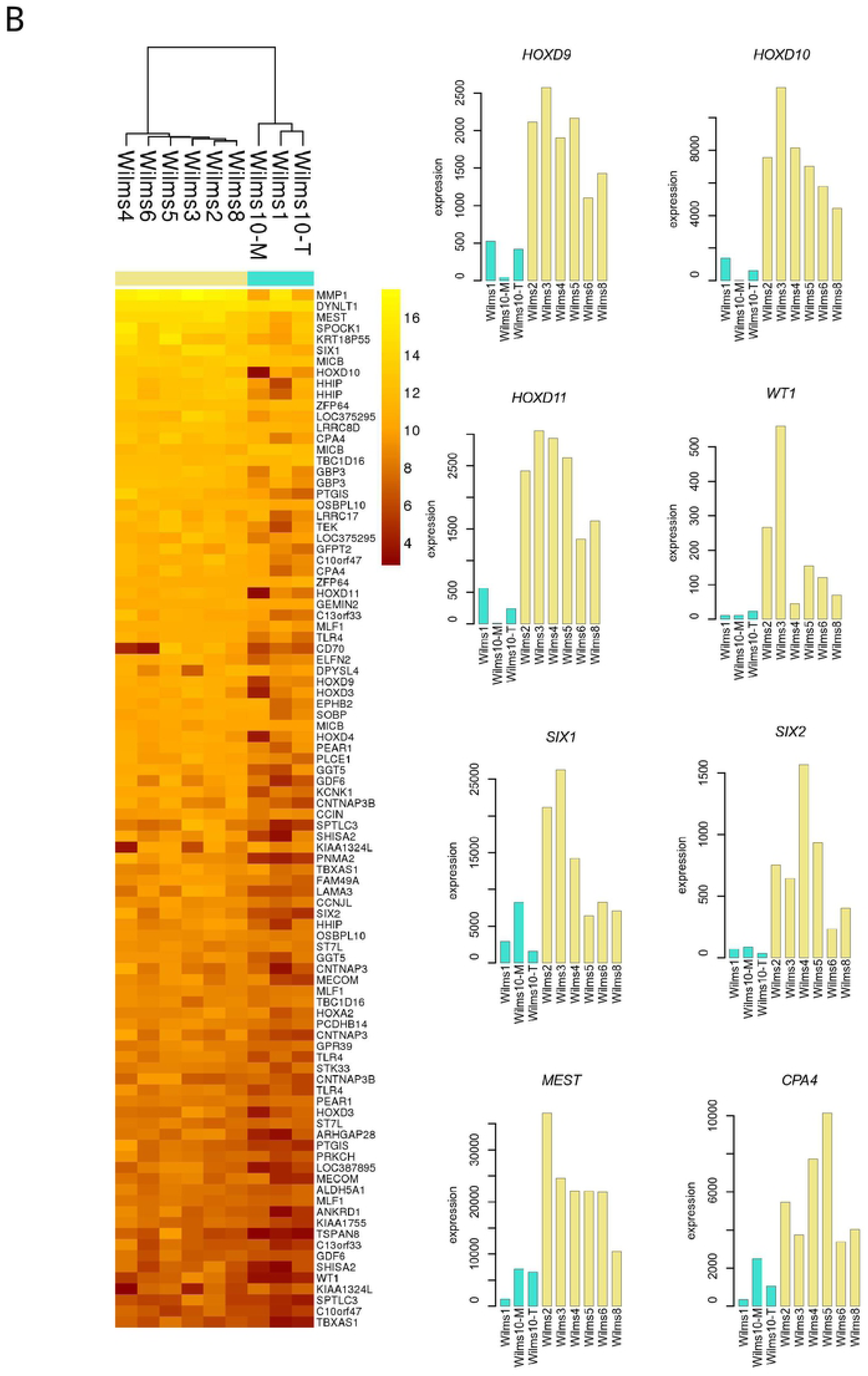
Differentially expressed genes in Wilms cells with and without *WT1* expression. **A)** Heat map of the genes expressed higher in cells without *WT1:* Wilms1, Wilms10T and Wilms10M (left) and expression level of selected genes with a higher expression in Wilms1, Wilms10T and Wilms10M (right). **B)** Heat map of genes expressed lower in cells without *WT1* (left) and expression of selected genes with a lower expression in Wilms cells without *WT1* expression (right).

To interrogate the biological function of the differentially expressed genes we performed a ToppGene analysis of the higher and lower expressed gernes and only for the genes with a lower expression a significant enrichment for GO terms is found. The first significant term is “embryonic skeletal system morphogenesis” with genes *HOXA2, HOXD3, HOXD4, HOXD9, HOXD10, HOXD11, WT1, SIX1* and *SIX2* all expressed at a lower level in the cells without *WT1.* Next are “embryonic organ development” and “anterior/posterior pattern specification” (Fig 11A). This shows that these biological processes are not active in these cells and that early embryonic patterning is erroneous. The specification of metanephric mesenchyme requires Hox and Wt1 proteins and as both are lacking this process might not be initiated. We also noticed that two genes with a lower expression, *MEST* and *CP4*, are both located in an imprinting cluster on chromosome 7q [42].

**Fig 11.**
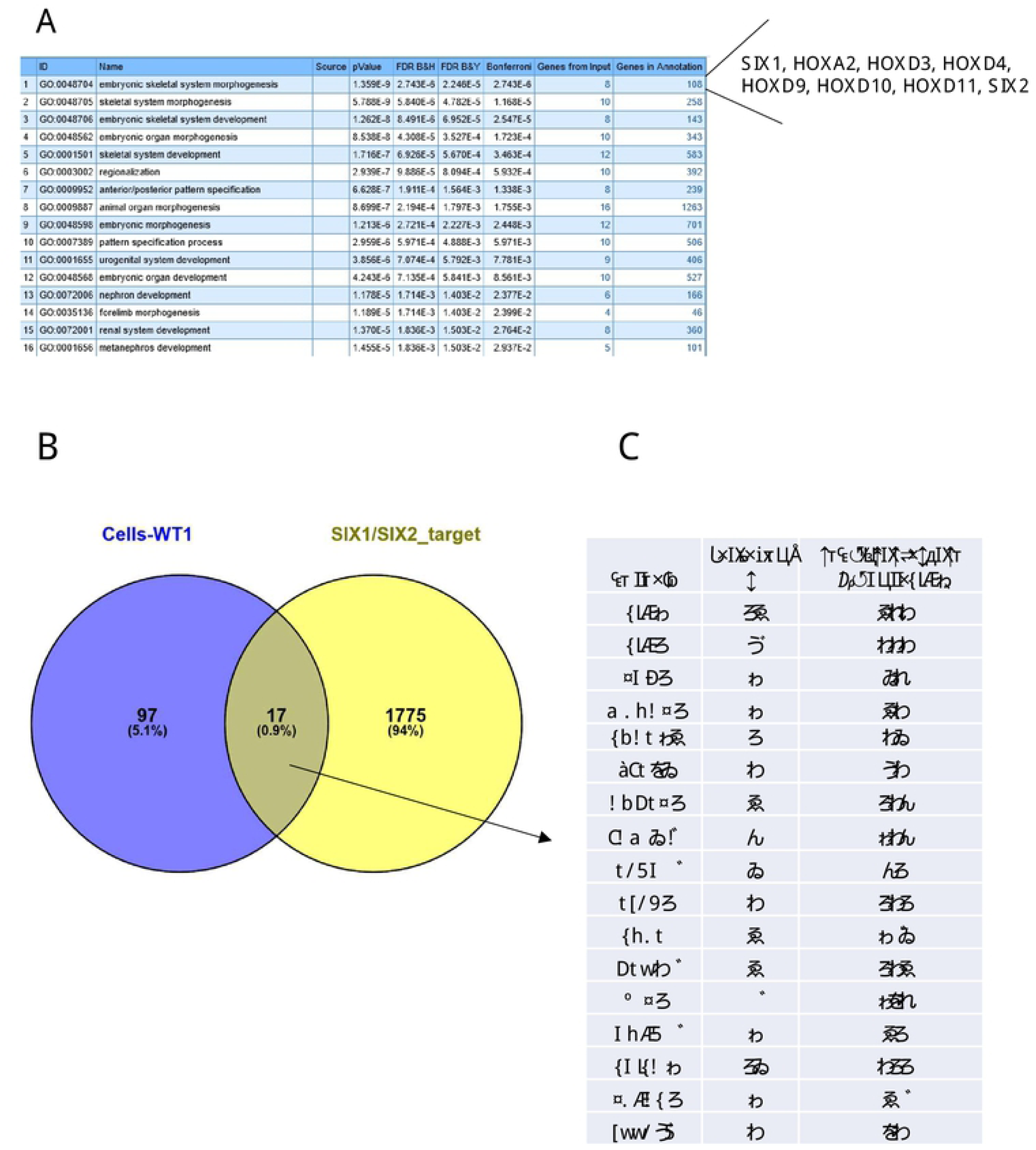
Analysis of differentially expressed genes in Wilms cells with and without mutant *WT1* expression. **A)** ToppGene result for the differentially expressed genes. **B**) Venny diagram shows that among the 114 differentially expressed genes in cells with and without *WT1* (blue circle) 17 are putative *SIX1/2* target genes (yellow circle, 1792 putative *SIX1/2* target genes identified by ÓBrien et al., 2016 [14]. **C)** A list of the 17 putative target genes with the number of peaks as detected with ChIP seq by ÓBrien et al., 2016 and the regulatory score that they have determined [14].

The lower expression of *SIX1* and *SIX2* was an interesting observation as these genes have a role in the balance of progenitor renewal and commitment. The disturbance of this balance could have a significant effect on tumor development. We therefore studied whether putative *SIX1* and/or *SIX2* target genes are among the differentially expressed genes. To address this question, we used the list of putative common *SIX1*/*SIX2* human target genes identified by ChIP-seq and expression studies in human and mouse kidney progenitors [14]. Indeed, 17 of the differentially expressed genes in the Wilms cell lines were identified as putative SIX1/2 target genes: *SIX2, SIX1, SISHA2*, *THY1, ZFP64, ANGPT1, FAM49A, PLCE1, SOBP, GPR39, WT1, HOXD9, MBOAT1, SOBP, LRRC8D, PCDH9* and *SNAP25* and the number of SIX binding sites and the regulatory scores are indicated in Fig 11B and C.

The correlation of no *WT1* expression with a low level of *SIX2* and *SIX1* might confirm that *WT1* is a major target for SIX2. In fact, multiple binding sites for Six2 were identified in the *Wt1* gene and therefore this gene is a highly likely and real Six2 target gene [14]. However, in the development of Wilms tumor the first step is haploinsufficiency for *WT1* expression either in the germline or during development in sporadic cases, followed by complete functional loss of both wild type alleles. Therefore, an alternative scenario could be that the complete loss of a functional wild type WT1 results in the reduced expression of *SIX2*, i.e., WT1 regulates *SIX2*. However, transfection of wildtype *WT1* in Wilms1 cells resulted in upregulation of *SIX1* not *SIX2* (data not shown). An alternative scenario is that the timing of *WT1* loss during nephrogenesis happens in a cell that has not turned on the expression of *SIX1/2* yet. If the loss of *WT1* occurs at a later time point when the cells have already activated the *SIX* genes this would result in tumors/cell lines with *SIX2* expression. Therefore, with these cell lines alone these alternative possibilities, loss of *SIX1/2* expression is due to loss of *WT1* or vice versa cannot be clarified.

### Comparison of Wilms tumor cell lines with human *WT1* mutant Wilms tumors and mouse *Wt1* mutant Wilms tumor models

Fukuzawa et al., 2017 analyzed *WT1* mutant human Wilms tumors in vivo for the expression of genes from the morphologically distinct cell types found in the kidney [43]. Their studies showed that these tumors contain cells with mesenchymal differentiation as well as UB like structures. Genes expressed in PT, RV, CSB, SSB, HL and DT were also identified but their expression was lower. Expression of *HOXD11*, a specific marker of MM was found in blastema as well as stromal cells, indicating that *WT1* mutant tumors are derived from the metanephric lineage [43]. *HOXD11* shows a low expression in cells with a complete lack of mutant *WT1* expression but a high expression in the other cell lines (Fig 10B). The cell lines that we have studied here have a similar expression pattern as cells from different compartments in tumors in vivo. This demonstrates that the expression of genes from all kidney cell types is indeed found in the same cells. Therefore, we confirm their conclusion from in vivo analyses of *WT1* mutant tumors that these are derived from stem cells with the capacity to differentiate into all cell types of the kidney. The more complete differentiation observed in vivo might be due to signaling between different cell types.

Two papers described Wilms tumor mouse models with *Wt1* ablation [44,45]. The first mouse model describes a somatic mosaic ablation of *Wt1* in a genetic background of biallelic expression of *Igf2* using a ubiquitously expressed, tamoxifen inducible transgene encoding Cre-recombinase [44]. The Wilms tumors that developed in these U-*Wt1-Igf2* animals had a triphasic histology, an early onset and these occurred at a high frequency. The cell of origin for the Wilms tumors in the U-*Wt1-Igf2* mouse was postulated to reside in the intermediate mesoderm. In the next approach they used *Cre* alleles that were expressed in *Foxd1* (F in stromal cells), *Cited1* (C in uninduced MM cells) or *Six2* (S in uninduced and induced MM) positive cells. In these settings they tested the effect of cell type specific ablation of *Wt1* in addition of either biallelic *Igf2* expression or coexpression with an activated β-Catenin protein through an exon 3 mutation (β-cat^S^) [45]. Their work showed that Wilms tumors developed only in mice when nephron but not stromal progenitors were targeted. Tumors were observed 1) when β-catenin was activated in nephron progenitors (NP cells) with *Six2* or *Cited1* promoters irrespective of *Wt1* ablation and 2) in C-*Wt1-Igf2* mice (targets only *Cited1^+^/Six2*^+^ cells) but not in S-*Wt1-Igf2* (targets *Cited1^+^/Six2*^+^ and *Six2* only cells [45]. The Wilms tumors that developed with *Wt1* ablation and biallelic expression of *Igf2* in NP cells (C-*Wt1-Igf2)* had a triphasic histology and they express early metanephric mesenchyme genes such as *Eya1, Osr1, Pax2,* and *Hoxa11* and postinduction genes. In contrast, when β-catenin is activated in NP cells (C-*Wt1*-*β-cat^S^* or S-*Wt1*-*β-cat^S^*) the tumors had an epithelial morphology and a low expression of renal mesenchyme genes, no expression of *Pax2* and expression of markers for committed NP cells and epithelial differentiation [45].

The genetics of the U-*Wt1-Igf2* mouse Wilms tumors (early onset) resembles that of *WT1* mutant human Wilms tumors. All *WT1* mutant human cell lines except Wilms4 from a WAGR patient, have a paternal duplication of chromosome 11p15, resulting in biallelic expression of *IGF2*. Furthermore, all except two cell lines have an additional mutant *CTNNB1* gene. Therefore, in most of these human Wilms tumor cells there are three genetic alterations: loss of wild type *WT1* in all, biallelic expression of *IGF2* and *CTNNB1* mutations, but none of the mouse tumor models harbor all three alterations. The human Wilms cell lines do not express *PAX2*, *EYA1* nor *CITED1,* but *OSR1, HOXA11, SIX2* and *FOXD1,* a combination not seen in the mouse tumor models.

Taken together these mouse models represent some aspects of *WT1* mutant human Wilms tumors and they show that the origin of Wilms tumors must lie in an early NP progenitor cell as we also show here for the human *WT1* mutant Wilms tumor cell lines. The work by Huang et al., revealed that a committed stromal cell cannot be the origin of Wilms tumors, but the high expression of interstitial genes that we observe in the Wilms tumor cell lines supports their close relationship to stromal cells and possibly a faulty differentiation into interstitial cell types as was postulated for *Pax2* negative mouse cells [6]. This further illuminates the origin of *WT1* mutant Wilms tumors in an early nephron progenitor with a disturbed differentiation pattern. The concomitant expression of early and postinduction NP genes indicates that they retain a multilineage potential.

### Expression of selected early kidney progenitor genes and their putative targets analyzed in Wilms cells

Crucial genes for NP cell pool maintenance in the mouse are *Wt1, Osr1, Eya1, Hox11* paralogs, *Six1* and *Six2.* In the cell lines only mutant *WT1* is present and *EYA1* is not expressed, whereas all other genes are positive. We searched publications describing mutant mouse phenotypes and putative targets/interactions partners of these genes and compared these to the data of Wilms cell lines.

*WT1-/-* embryos express *Pax2* and *Gdnf* mRNA prior to aberrant apoptosis [19] and the *WT1* negative Wilms tumor cells do not express *PAX2*, but *GDNF. Osr1* acts upstream of *Pax2, Sall1, Eya1, Six2* and *Gdnf* during kidney development and *Osr1* mutant embryos lack the expression of these genes [46]. The Wilms tumor cells express variable levels of *OSR1* and *SIX2* (Fig 10B). *SIX1, SALL1* and *GDNF* are all robustly expressed but *PAX2* is undetectable. In the mouse *Pax2* and *Eya1* are needed for the expression of *Gdnf*. In contrast, in the Wilms cells *GDNF* is expressed in the absence of *EYA1 and PAX2*, suggesting an alternative mechanism for activation in the human tumor cells.

In *Pax2* mutant cap mesenchyme there is no expression of *Cited1, Ncad (CDH2)* and at some stage coexpression of *Six2* and *Foxd1* was observed. In the *PAX2* negative Wilms cells, *SIX2, FOXD1* and *CDH2* are expressed, whereas *CITED1* is negative. Furthermore, in mice the expression of *Eya1, Sall1* and *Six1* is *Pax2* independent and in the human Wilms tumor cells *SIX1* is highly expressed and *SALL1* at a lower level, confirming their *PAX2* independent activation.

In mice the Hox11 paralogs complex with Eya1 and Pax2 to drive expression of *Six2* and *Gdnf* [47]. In the Wilms cell lines *HOXD11* is variably expressed and *HOXA11* at a lower level. As both *EYA1* and *PAX2* are not detectable in Wilms cells a [Hoxa11-Pax2-Eya1] complex cannot be formed. However, the cells express *EYA2, PAX3* and *PAX8*, suggesting that alternative complexes might be formed with other family members to regulate the expression of *GDNF* and *SIX2*.

A loss-of function for *Six1* and *Six2* genes results in kidney agenesis. In the *Six1* mutant, the levels of *Six2, Pax2* and *Sall1* are reduced, whereas *Eya1* was unaffected and expression of *Wt1* is normal [48]. This indicates that *Six1* is upstream of *Six2*. In the Wilms cells *SIX1* is highly expressed, *SALL1* lower, *SIX2* is variable and *PAX2* is lacking. In summary, this short survey of mouse data compared to the human Wilms tumor cells unravels that different mechanism must exist in the human tumor cells to control these genes.

## Summary and conclusion

Our analyses uncovered that the Wilms tumor cells coexpress genes from all kidney compartments and genes from more mature cells. As all genes are expressed in the same cells, this confirms that the cell of origin for *WT1* mutant Wilms tumors is a common precursor for all different kidney cell types. However, a faulty concomitant differentiation in all lineages has been initiated. In the precursor cells that lack wild type *WT1,* important factors for a normal nephrogenic differentiation program are missing, e.g., *CITED1, PAX2, EYA1* and *MEOX1* while other NP genes are expressed e.g., *SIX2, OSR1*, *SIX1* and *SALL1*. Although most of these cell lines have an activated Wnt signalling pathway that normally induces differentiation, this does not occur in the Wilms cell lines. Furthermore, the analysis of coexpression of specific genes with their putative targets shows that in these cells other gene regulations are operative. To further explore these developmental gene regulations, the Wilms tumor cell lines can be used as model systems. The function of these genes can be studied by transfection and analysis of the outcomes. With the CRISPR-Cas9 technique the mutant *WT1* gene can be replaced by a wild type *WT1* as well the mutant *CTNNB1* gene with a wild type gene. Using such manipulated cells in a kidney organ system should reveal their differentiation potential into the various cell types of the kidney and determine whether they are true kidney stem cells.

## Supporting information

**S1 Fig. Overlapping genes from the normal human nephrogenic clusters 4A, 4B, 4C and 4D [32].** Cluster 4A, corresponds to 222 genes, 209 are exclusively expressed; cluster 4B, corresponds to 59 genes, 47 are exclusive; cluster 4C corresponds to 271 genes and 217 are exclusive; cluster 4D corresponds to 1380 genes and 1311 are exclusive

**S2 Fig. *KeyGenes* expressed in Wilms cells corresponding to the different kidney clusters [38].** Heat maps of the expressed *KeyGenes* in Wilms cell with lists of all marker genes from each normal human cluster below the maps [38]. If a gene is also expressed in another cluster this is listed in the same line. Expression of genes in Wilms cell lines of all clusters are shown as heat maps. Symbols from BACs e.g., RP11-609N14.1 were not included in the calculations as no genes were represented by these.

**S3 Fig. Heat map of expression of literature genes [38] in the Wilms cell lines.** One gene, *GDNF* is present in two clusters, NPC and IPC, but only listed in NPC.

**S4 Fig. Highest expressed genes in the Wilms8 cell line and classification to compartments.** Here the number of genes with a Z-score expression in normal human fetal kidney between 3 and 4 (Semrau database) are shown that are highly expressed in Wilms8. The different colors indicate the expression level.

**S5 Fig. Genes expressed higher in group2 cell lines and their classification to normal kidney compartments.** Here the genes with a p-value <0.05, expression >200 and a ratio of >1.33 were analyzed for their allocation to normal human fetal kidney compartments. The result shows the expression of these genes in normal kidney, not in the Wilms cell lines (Semrau database).

**S6 Fig. Genes expressed lower in group 2 cell lines and their classification to normal kidney compartments.** The genes with a significant lower expression in group 2 (p<0.05, expression >200 and a ratio of <0,75) were studied for their expression in normal human fetal kidney (Semrau database).

**S7 Fig. Expression of genes from the Wang clusters [15] in Wilms cell lines. A)** percentage of genes from each compartment expressed in Wilms cell lines, cut off >1000 in any of the cell lines. **B)** The numbers of genes in each compartment and percentages of expressed genes in Wilms cell lines.

**S1 Table (docx):** Key genes for each Lindström cluster and highest expressed genes in Wilms cell lines

**S2 Table (xls):** Genes expressed in Wilms cell lines (<1000) from all Lindström clusters

**S3 Table (xls):** Genes expressed in Wilms cell lines (<1000) from all Hochane clusters

**S4 Table (docx):** Marker genes for each Hochane cluster and highest expressed genes in Wilms cell lines

**S5 Table (xls):** Genes expressed in Wilms cell lines (<1000) from all Wang clusters

**S6 Table (docx):** Marker genes from Wang clusters and highest expressed genes in Wilms cell lines

## Author contributions

**Conceptualization:** B. Royer-Pokora

**Data Curation:** B. Royer-Pokora, M. Beier

**Formal Analysis:** W. Wruck, M. Beier

**Investigation:** B. Royer-Pokora, W. Wruck, M. Beier

**Methodology:** B. Royer-Pokora

**Software:** M. Beier

**Visualization:** B. Royer-Pokora, M. Beier

**Writing – Original Draft Preparation:** B. Royer-Pokora

**Writing – Review & Editing:** B. Royer-Pokora, W. Wruck, M. Beier

